# Neonatal Amygdala Mean Diffusivity: A Potential Predictor of Emotional Face Perception

**DOI:** 10.1101/2024.02.04.578788

**Authors:** Niloofar Hashempour, Jetro J. Tuulari, Harri Merisaari, John D. Lewis, Tuomo Häikiö, Noora M. Scheinin, Saara Nolvi, Riikka Korja, Linnea Karlsson, Hasse Karlsson, Eeva-Leena Kataja

## Abstract

The ability to differentiate between different facial expressions is an important part of human social and emotional development that begins in infancy. Studies have shown that within the first year of life, infants develop a distinctive attentional bias towards fearful facial expressions. Investigations into the neural basis for this bias have highlighted the significance of the amygdala. The amygdala’s role in directing attention towards fearful facial expressions underscores its importance in early emotional development, significantly influencing how infants interpret and react to facial expressions. To date, no studies have been conducted to investigate the associations between the amygdala microstructure and infants’ perception of emotional faces. This study aimed to elucidate this relationship while also investigating whether this association is sex specific. We measured the amygdala microstructural properties using diffusion tensor imaging mean diffusivity (MD) measurements in 40 healthy infants aged 2 to 5 weeks. Eye tracking was used to assess attention disengagement from fearful vs. non-fearful (happy and neutral) facial expressions as well as scrambled non-face control picture at 8 months. Generally, infants were age-typically less likely to disengage from fearful faces than from non-fearful faces towards salient distractors. A significant negative association was observed between the right amygdala MD measures and disengagement probability from fearful faces in the overall sample. Moreover, there was a positive association between the bilateral amygdala MD measures and the disengagement probability from scrambled non-face control picture in girls. These results indicate that the amygdala MD is associated with attention disengagement processes already in infancy, both in fear processing and in non-emotional conditions. Specifically, these findings highlight the role of the amygdala microstructure in modulating attentional processes, which may have implications for emotional regulation and susceptibility to emotional dysregulation later in life.

## Introduction

Attentional preference for social stimuli such as faces is an important prerequisite for humans’ social and emotional development (Prunty et al., 2020; Reynolds & Roth, 2018). Newborns typically demonstrate a predilection for faces and tend to sustain their visual attention on them over non-social stimuli immediately after birth, and this trait is tractable throughout infancy (Farroni et al., 2005; Johnson et al., 1991; Libertus et al., 2017). Infants as young as 3 months old have been shown to have a preference for faces over other objects (Farroni et al., 2005), and this preference becomes even stronger by 6 months of age (Kelly et al., 2019). In addition to a preference for faces, there is also evidence that infants can distinguish between different types of facial expressions. Studies have shown that infants exhibit an attentional bias towards fearful facial expressions by the age of 7 months compared to other expressions, such as happy faces (C. A. Nelson & Dolgin, 1985; Peltola et al., 2009). Additionally, at 7 months, infants take longer to shift their attention away from fearful faces compared to happy or neutral faces when a salient distractor stimulus is presented to the left or right side of the face (Peltola et al., 2008, 2009, 2013). Moreover, compared to happy or neutral faces, 12-month-old infants focused their visual attention on fearful faces significantly more often (Nakagawa & Sukigara, 2012). While the cause of infants having an attentional bias towards fearful faces has not been completely established, it has been suggested that it may stem from the distinctiveness of fear expressions, as in the infant’s environment, fear is not encountered as frequently as other emotions (C. A. Nelson & Dolgin, 1985) and some particular features of fearful faces, such as large, open eyes (Peltola et al., 2008). This attentional bias toward fearful faces may also influence later social skills and social behaviors (Charles A. Nelson, 2001), and individual differences in the magnitude of early fear bias may precede individual variance in fear sensitivity and differences in empathy and anxiety proneness (Marsh, 2016).

The exact neural processes that drive infants’ attention to fearful faces are not yet clear. It is suggested that the development of brain structures in the limbic system, specifically the amygdala, plays a role in infants’ attentional processes, and the processing of fearful faces (Tuulari et al., 2020; Vuilleumier, 2005; Whalen et al., 2001). Studies suggest that the amygdala is crucial for threat detection, as evidenced by difficulties in processing fearful facial expressions among individuals with amygdala lesions. (Adolphs et al., 1999; Whalen et al., 2001). Amygdala development begins during the embryonic period and continues throughout infancy (Humphrey, 1968; Uematsu et al., 2012). The amygdala is one of the main centers for facilitating an individual’s adaptation to their environment (Cho et al., 2013). Therefore, it processes emotional stimuli efficiently and rapidly (Balderston et al., 2014). Various sensory modalities, such as auditory, visual (Morrow et al., 2019), and olfactory stimuli, provide information to the amygdala (Baysal Akin & Onat, 1995). Then, the amygdala facilitates the right behavioral response by activating the hypothalamic-pituitary-adrenal axis and releasing stress hormones (Smith & Wylie, 2006) or by directly stimulating autonomic and behavioral reactions (Watson et al., 2010). The amygdala responds to all emotional facial expressions (Fitzgerald et al., 2006), but expressions of fear elicit a stronger response than those of other emotions (Morris et al., 1996; Tuulari et al., 2020). This evolutionary adaptation provides a significant survival advantage in situations when a potential threat needs to be recognized (Adolphs, 2010; Susskind et al., 2008). Neuroimaging studies in infants have linked amygdala volume and diffusion properties to conditions like prenatal maternal distress, which are commonly associated with fearful dispositions (Acosta et al., 2020; Favaro et al., 2015; S. J. Lehtola et al., 2020; Rifkin-Graboi et al., 2013; Wen et al., 2017). While one study using a partially overlapping sample with the current study, reported a link between amygdala volumes and infants’ attentional bias toward fearful facial expressions (Tuulari et al., 2020), no studies have examined the associations between amygdala diffusion properties and perception of fearful faces. Examining this relationship may provide insights into how the amygdala’s microstructure is related to the neural basis of fear development at an early age, as well as its long-term implications.

Diffusion tensor imaging (DTI) can serve as an intermediate phenotype to study the associations between the microstructure of the amygdala and emotional facial processing. DTI measures such as mean diffusivity (MD) reflect the average rate of diffusion of water molecules, regardless of their directionality (Stebbins, 2010). Higher amygdala MD is found in patients with anxiety (Juranek et al., 2012), impaired cognitive performance, and neurovegetative diseases like Alzheimer’s (Cherubini et al., 2010). Increased MD measures are linked with shrinkage of neurons, and synapse loss, as well as immaturity and tissue degeneration (Cherubini et al., 2010; De Gennaro et al., 2011; Gillespie et al., 2017; Kantarci et al., 2005). MD generally decreases throughout childhood and adolescence in healthy individuals and is associated with higher cognitive performance (Takeuchi et al., 2016; Tamnes et al., 2010; Yoshida et al., 2013). These findings suggest that higher MD measures in the amygdala may be a common feature across various psychiatric conditions and fear (dys)regulation, indicating a shared neurobiological vulnerability.

In the current study, we examined how the newborn MD of the amygdala related to the attentional bias towards different facial expressions at 8 months. Infants at 8 months, typically show a high preference for fearful vs. non-fearful facial expression (neutral and happy), but variance also exists in this bias, making it a suitable measure for attention bias studies. Further, to see if these connections are sex-specific, we explored the sexually dimorphic associations of facial expressions with the amygdala MD measures. This was an exploratory study, and no hypothesis for the association or direction of the effect was set. We note that studies investigating the diffusion properties of the amygdala in associations with emotional processes in infancy are lacking.

## Methods

### Participants

The study included N = 40 infants (50% males), enrolled in the FinnBrain Birth Cohort Study (Karlsson et al., 2018). Pregnant mothers were recruited to the study at their first ultrasound visit during gestational week (gwk) 12 at three maternal well-fare clinics performing ultrasound scans for the women giving birth at Turku University Hospital in the Southwest Finland Hospital District and the Åland Islands in Finland between December 2011 and April 2015. The background information was collected with questionnaires at gwk 14 and/or 34 (Table 1). Obstetric data were retrieved from national birth registries (National Institute for Health and Welfare, www.thl.fi) (Table 1). The inclusion criteria for MRI scans were birth weights >2500 g and gestational ages ≥37 weeks. Exclusion criteria were previously diagnosed congenital abnormalities of the central nervous system (CNS) and abnormal findings in a prior magnetic resonance imaging (MRI) scan. All infants with a completed MRI at 2–5 weeks after birth and a valid eye-tracking assessment at 8 months were included in the current analyses. The eye-tracking assessments and imaging were conducted as part of the visits of the FinnBrain Child Development and Parental Functioning Lab and Neuroimaging Lab, respectively, at the University of Turku.

**Table 1.**
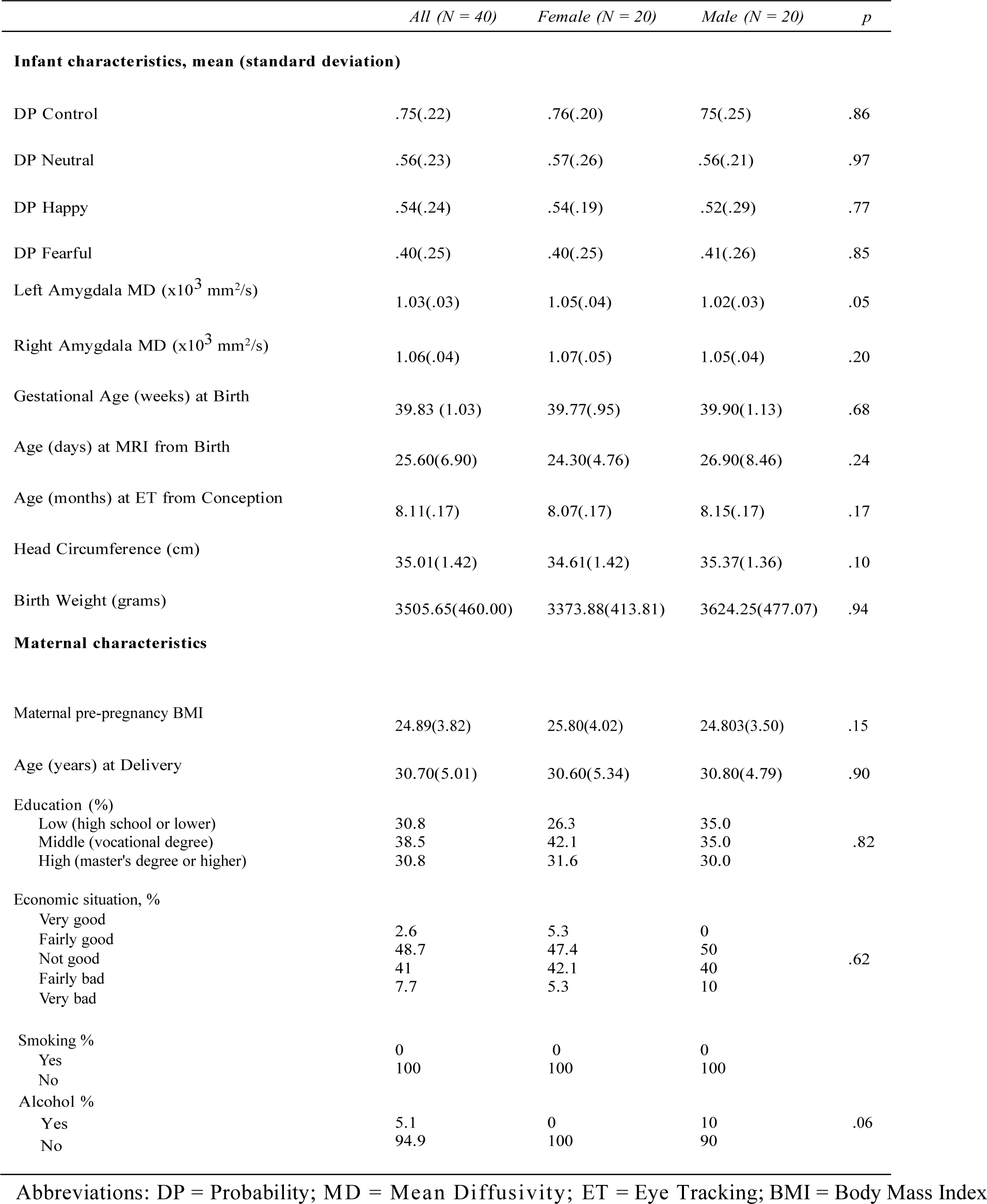
The sample characteristics.

Written informed consent was obtained from all parents. This study was conducted in accordance with the Declaration of Helsinki and was approved by the Ethics Committee of the Hospital District of Southwest Finland (ETMK: 31/180/2011).

### Diffusion-weighted imaging data acquisition and processing

MRI scans were performed using the Siemens Magnetom Verio 3T scanner (Siemens Medical Solutions, Erlangen, Germany). Please see a description of the details of the scanning protocol in our previous publication (Lehtola et al., 2019). We used a 96-direction diffusion-weighted imaging (DWI) protocol, which was divided into three parts (composed of 31, 32, and 33 non-identical diffusion encoding directions). The diffusion encoding directions were evenly distributed across the three-dimensional space in the full 96-direction protocol in each part (each part with a duration of approximately 6 min). For this study, 60 diffusion directions with b = 1000 s/mm2 and three b = 0 s/mm2 images were chosen to provide better tensor model fitting and outcomes. The sequences were acquired using the Spin Echo-Echo Planar Imaging sequence at 2 mm^3^ isotropic resolution (FOV 208 mm; 64 slices; TR 8500 ms; TE 90 ms). DWI data were pre-processed as previously described in detail in (Merisaari et al., 2019), summarized here for sake of brevity. DTIprep and visual quality control were used to censor bad diffusion directions, eddy/motion correction was applied with FSL tools, with rotation of B-matrix. To yield MD measures, we processed the 4D diffusion dataset with FSL’s dtifit, using a brain mask obtained with BET brain extraction tool on an average b= 0 s/mm2 image of pre-processed DWI, to limit the modeling exclusively to brain tissue.

### Labeling the structures

The left and right amygdala were identified in each subject using label-fusion-based methods. These methods depend on achieving good registration between the entries using constructed library templates, based on each study subject. To ensure good registrations, we constructed a population-specific template, and a library of templates that represented the morphological variation in the sample. We constructed a population-specific base template based on the methods described in Fonov et al (Fonov et al., 2011), warped that template to the 21 subjects that best represented the morphological variation in the whole sample, and manually labeled the amygdala on each template as previously described (Hashempour et al., 2019). We then unwarped these 21 manual segmentations, created consensus segmentation labels on the base template via a voxel-wise majority vote, and constructed a library of warped versions of the labeled template that represented the morphological variation in the sample for those structures. This library was then used to label the individual brains via label-fusion-based labeling methods. A detailed description of methods can be found in Acosta et al. (Acosta et al., 2020).

### Extracting MD measures

The diffusion data were registered with the T1-weighted data, and the DTI measure extracted for the structures of interest as follows: 1) The B0 volume, which contains raw T2 signal devoid of diffusion weighting, was extracted from the diffusion datasets. Subsequently, antsRegistration was employed to perform a rigid registration, aligning the B0 volume with the nonuniformity-corrected T1-weighted data in native space; 2) The resultant transform was then concatenated with the transform from native to stereotaxic group average template space, and the result was applied to each of the DTI maps to overlay them on the T1-weighted data in stereotaxic space in a 0.5 mm isotropic voxel. For each DTI map, the MD measures within the amygdala were calculated, taking into account the partial volume effects on the borders of each structure. The resolution of the diffusion data was 2 mm isotropic, thus, complete elimination of partial volume effects would have discarded a large portion of the measures within small structures such as the amygdala. To eliminate most of the partial volume effects while also retaining a sufficient set of DTI measures for the statistical analysis, we eroded the amygdala labels by 1.5 mm before using the eroded labels as masks for the calculation of the mean of the remaining measures in each of the DTI maps.

### Eye-tracking-based assessment of infant attention disengagement from emotional facial expressions

Eye tracking (EyeLink1000+, SR Research Ltd, Toronto, Ontario, Canada) and an overlap paradigm (Peltola et al., 2008) were used to study infant attention disengagement from a centrally presented face or a scrambled non-face control picture to a salient lateral distractor. This paradigm has been used in several previous studies to examine infants’ attentional biases for faces vs. non-faces (Peltola et al., 2008) and specifically for fearful vs. non-fearful facial expressions (neutral and happy), (Nakagawa & Sukigara, 2012; Peltola et al., 2009; Tuulari et al., 2020), and we have reported the measurement details in our previous papers from the same sample (Tuulari et al., 2020).

In the overlap paradigm, the infants were shown photographs of two different models portraying happy, fearful, and neutral faces together with scrambled non-face control picture (Peltola et al., 2008). Altogether, a set of 48 trials were presented, including 12 trials per condition (each emotion and the scrambled non-face control picture) and comprising 18 photographs of each model, and 12 scrambled non-face control pictures, in a semi-random order. During the experiment, the infants were first shown a picture of a face (or a scrambled non-face control picture) in the center of the screen for 1000 ms (Figure 1). Then, a lateral distractor (black and white checkerboard or circles) appeared on either the left or right side of the face (a visual angle of 13.6◦) for 3000 ms, simultaneously with the face. One trial lasted for 4000 ms. The order of the central stimuli was semi-randomized, with the constraint that the same stimulus not be presented more than three times in a row. The lateral stimulus was selected and presented randomly for each trial.

**Figure 1.**
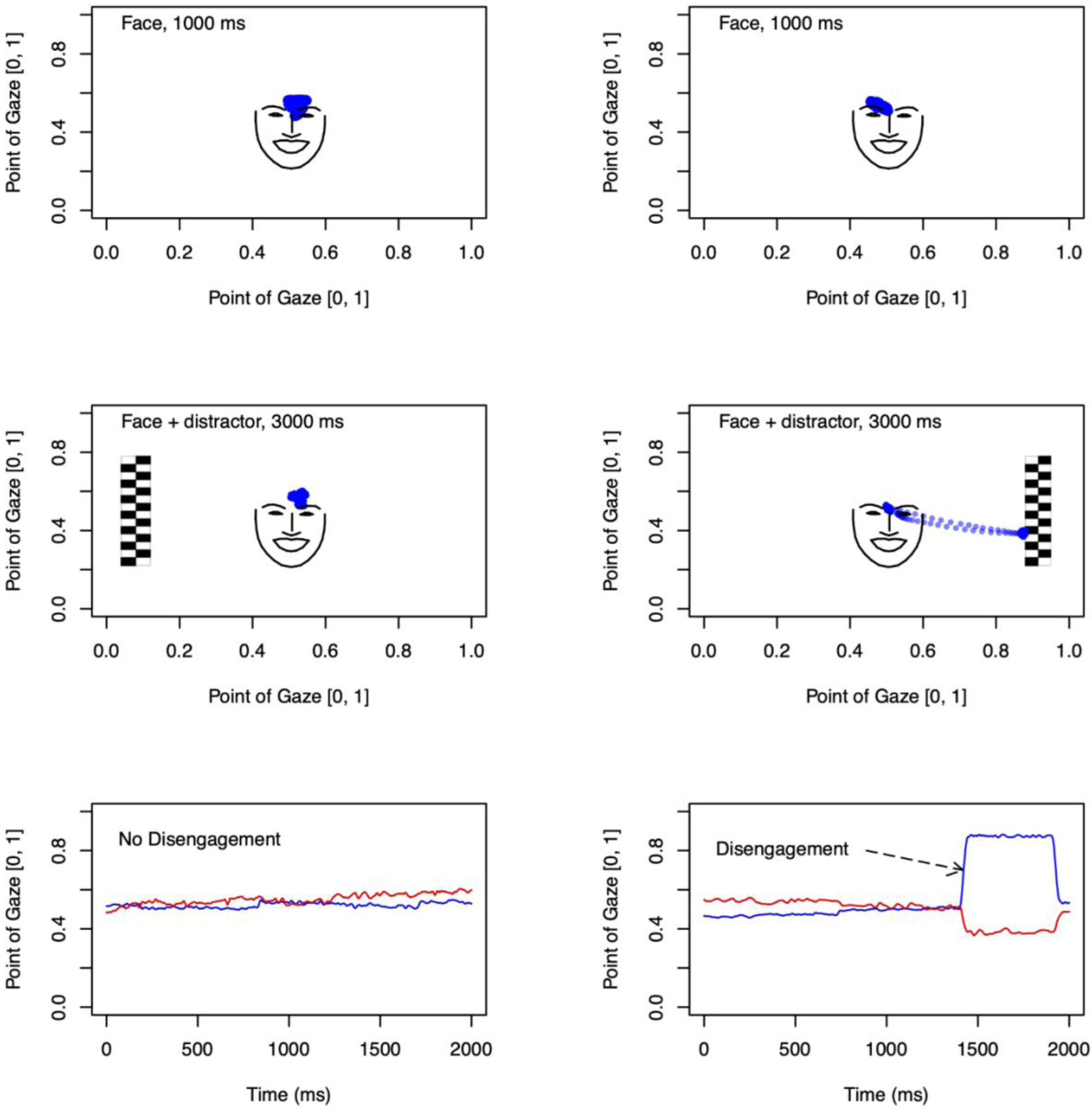
On the left, an example of a no disengagement trial, and on the right, an example of a trial with disengagement. The upper panel shows fixation locations (blue circles) within the first 1000 ms of the trial with just the non-face stimulus. The middle panel shows fixation locations after the distractor has appeared until the end of the trial. The lower panel shows the x (blue) and y (red) positions of the eye relative to the screen as a function of time (first 2000 ms of the trial only).

### Extraction of disengagement probabilities

The trial data, comprised of timestamps for the onset times of central and lateral pictures and the xy coordinates of the participants’ gaze position (500 samples/second), were stored as text files, and analyzed offline using a library of Matlab 2016b (Mathworks, Natick, MA) scripts for preprocessing and analysis of saccadic reaction times from eye-tracking data (Leppänen et al., 2015). Partially incomplete eye tracking data is common, and we used the following quality control criteria based on prior studies (Leppänen et al., 2015) to retain trials for the analysis. First, the infant’s gaze had to stay in the area of the central stimulus for >70 % of the time preceding gaze disengagement or the end of the analysis period. Second, trials had to have a sufficient number of valid samples in the gaze data (no gaps or missing eye position data >200 ms). Thirdly, trials had to have valid information about the eye movement from the central to the lateral stimulus (i.e., the eye movement did not occur during a period of missing gaze data). In order to be included in the analyses, the infant had to provide ≥3 valid trials for each stimulus condition. On average, the infants had 9.5, 9.7, 9.7, and 10.2 valid trials in the non-fearful (neutral and happy), fearful, and scrambled non-face control picture respectively. In these trials, the mean probability of attention disengagement (DP) from the central to the lateral stimulus was calculated for each stimulus condition and used in statistical analyses.

### Statistical analyses

Statistical analyses were performed using R version 4.1.1 (R Core Team, 2016). First, multiple linear regression was used to investigate the associations between the right and left amygdala MD measures and the disengagement probability in non-fearful faces (neutral and. happy), fearful faces and scrambled non-face control picture, while controlling for infant age from birth at the time of the MRI scan (MD ∼ disengagement probability + covariate). Second, sex-specific associations were investigated by including an interaction between infant sex and disengagement probability from faces (MD ∼ disengagement probability*sex + covariate). In all analyses, the level of statistical significance was set at p < .05. Given the exploratory nature of the study, no correction for multiple testing was carried out.

## Results

### Potential confounders influencing the association between the amygdala MD and the disengagement probability

There was a correlation between the left and right amygdala MD measures (r = .70, p *=* .001). The disengagement probability from the scrambled non-face control picture was negatively correlated with infant birth weight (r = -.38, p *=* .019). A negative correlation between the left amygdala MD and age at the MRI date was observed (r =-.41, p *=* .007). Age at eye-tracking date was negatively correlated with disengagement probability from the scrambled non-face control picture (r =-.37, p *=* .024) and positively with infant birth weight (r =-.34, p *=* .042). The intercorrelations between the variables are presented in Table 2.

**Table 2.**
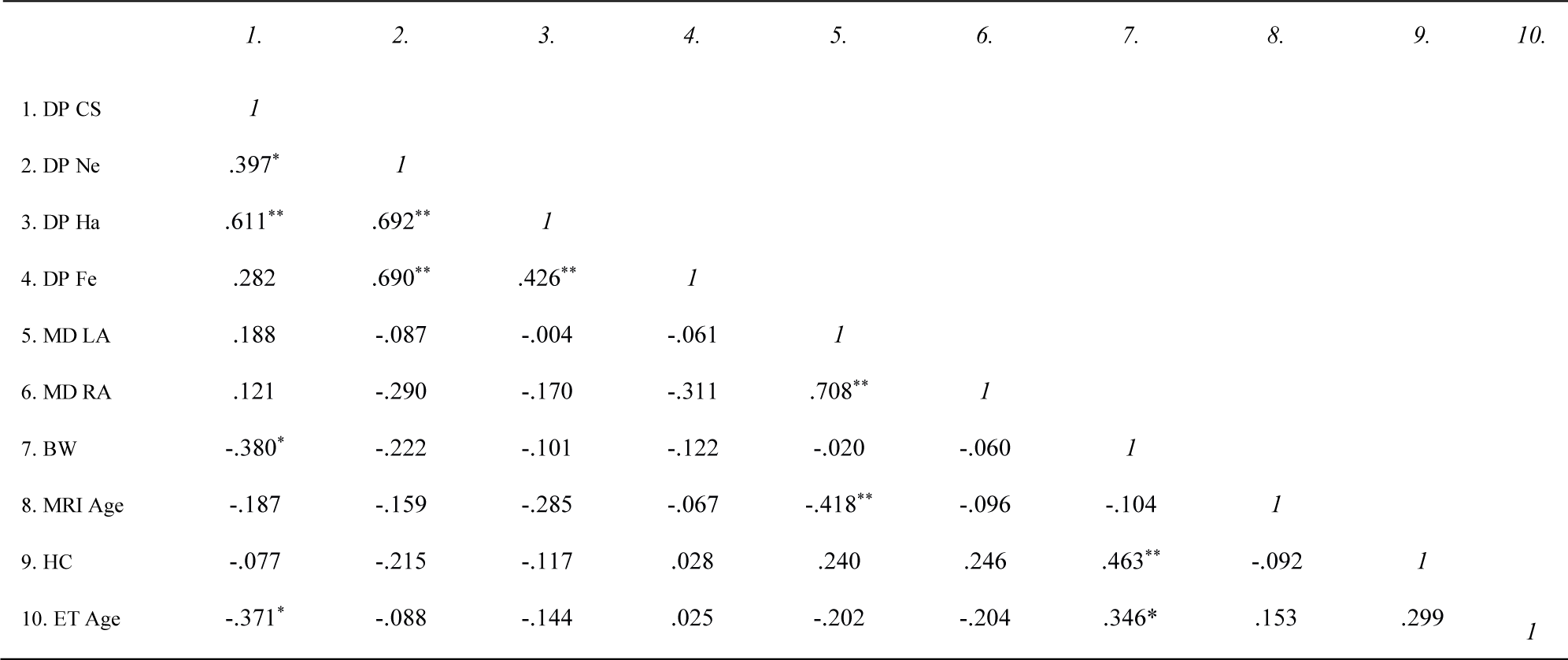
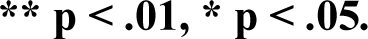
The Pearson correlations for disengagement probability (DP) from scrambled non-face control picture (Cs) neutral (Ne), happy (Ha), and fearful (Fe) stimuli; MD measures of the left (MD LA) and right amygdala (MD RA); birth weight (BW); age at the MRI date (MRI Age); head circumference (HC) and age at eye tracking (ET Age).

|  | 1. | 2. | 3. | 4. | 5. | 6. | 7. | 8. | 9. | 10. |
| --- | --- | --- | --- | --- | --- | --- | --- | --- | --- | --- |
| 1. DP CS | *1* |  |  |  |  |  |  |  |  |  |
| 2. DP Ne | .397* | *1* |  |  |  |  |  |  |  |  |
| 3. DP Ha | .611** | .692** | *1* |  |  |  |  |  |  |  |
| 4. DP Fe | .282 | .690** | .426** | *1* |  |  |  |  |  |  |
| 5. MD LA | .188 | -.087 | -.004 | -.061 | *1* |  |  |  |  |  |
| 6. MD RA | .121 | -.290 | -.170 | -.311 | .708** | *1* |  |  |  |  |
| 7. BW | -.380* | -.222 | -.101 | -.122 | -.020 | -.060 | *1* |  |  |  |
| 8. MRI Age | -.187 | -.159 | -.285 | -.067 | -.418** | -.096 | -.104 | *1* |  |  |
| 9. HC | -.077 | -.215 | -.117 | .028 | .240 | .246 | .463** | -.092 | *1* |  |
| 10. ET Age | -.371* | -.088 | -.144 | .025 | -.202 | -.204 | .346* | .153 | .299 | *1* |

**\*\* $p < .01$ , \* $p < .05$ .**

### The associations between the amygdala MD measures and the disengagement probability from emotional facial expressions

The infants were less likely to disengage from fearful (the mean probability of attention disengagement DP, *M* = .40) than happy (*M* = .53), neutral (*M* = .56), or scrambled non-face control picture (*M* = .75) faces, demonstrating an age-typical bias for fear. In the overall sample, a significant association was observed between the right amygdala MD measures and the disengagement probability from the fearful face after controlling for infant age at the time of MRI scan. The higher the right amygdala MD measures, the lower the probability to disengage from the fearful face (*ß* = -.058, *p =* .047) (Figure 2) (Table 3). No significant associations were found between the MD measures in either the left or right amygdala and DPs in other conditions (*p > .05*).

**Figure 2.**
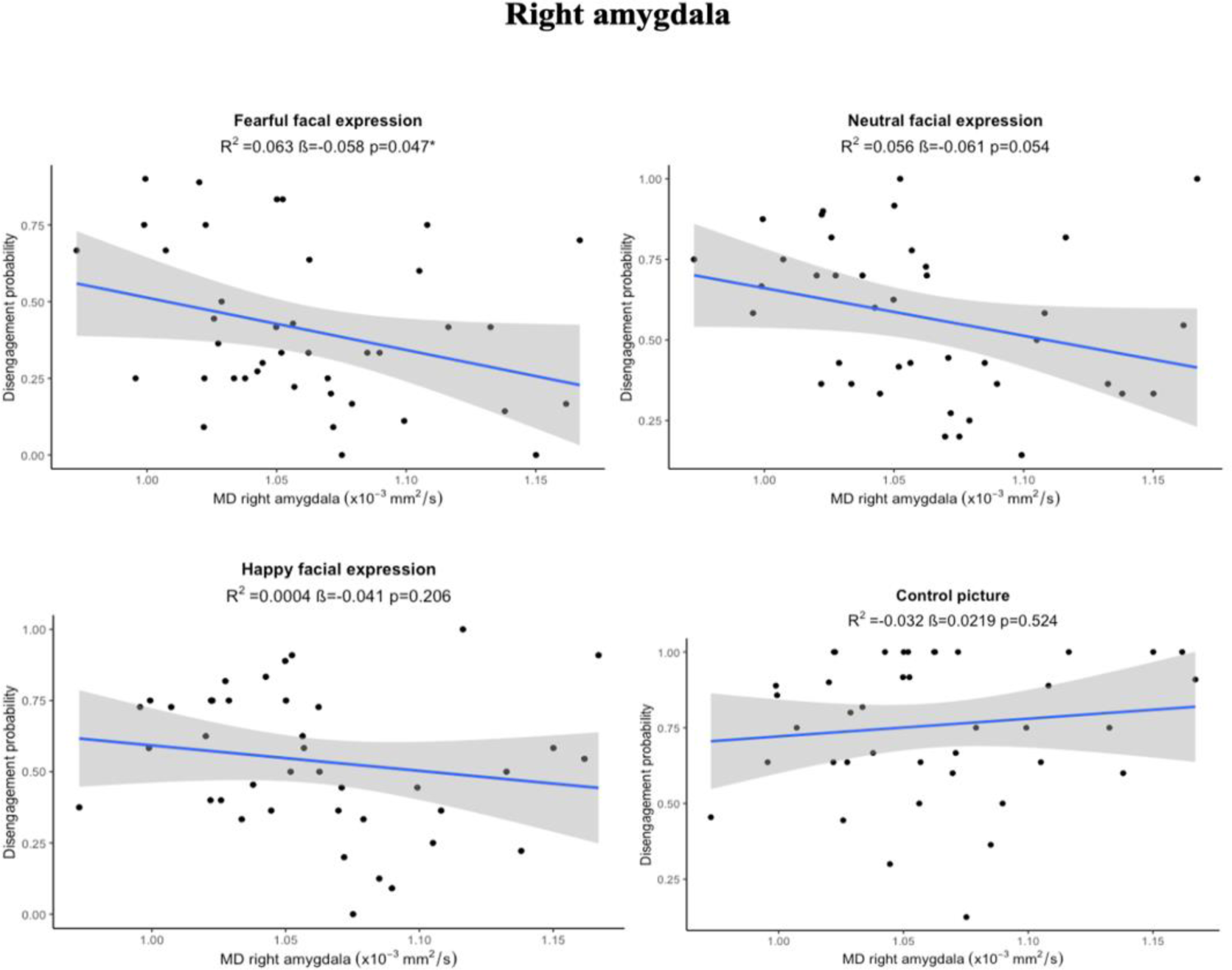
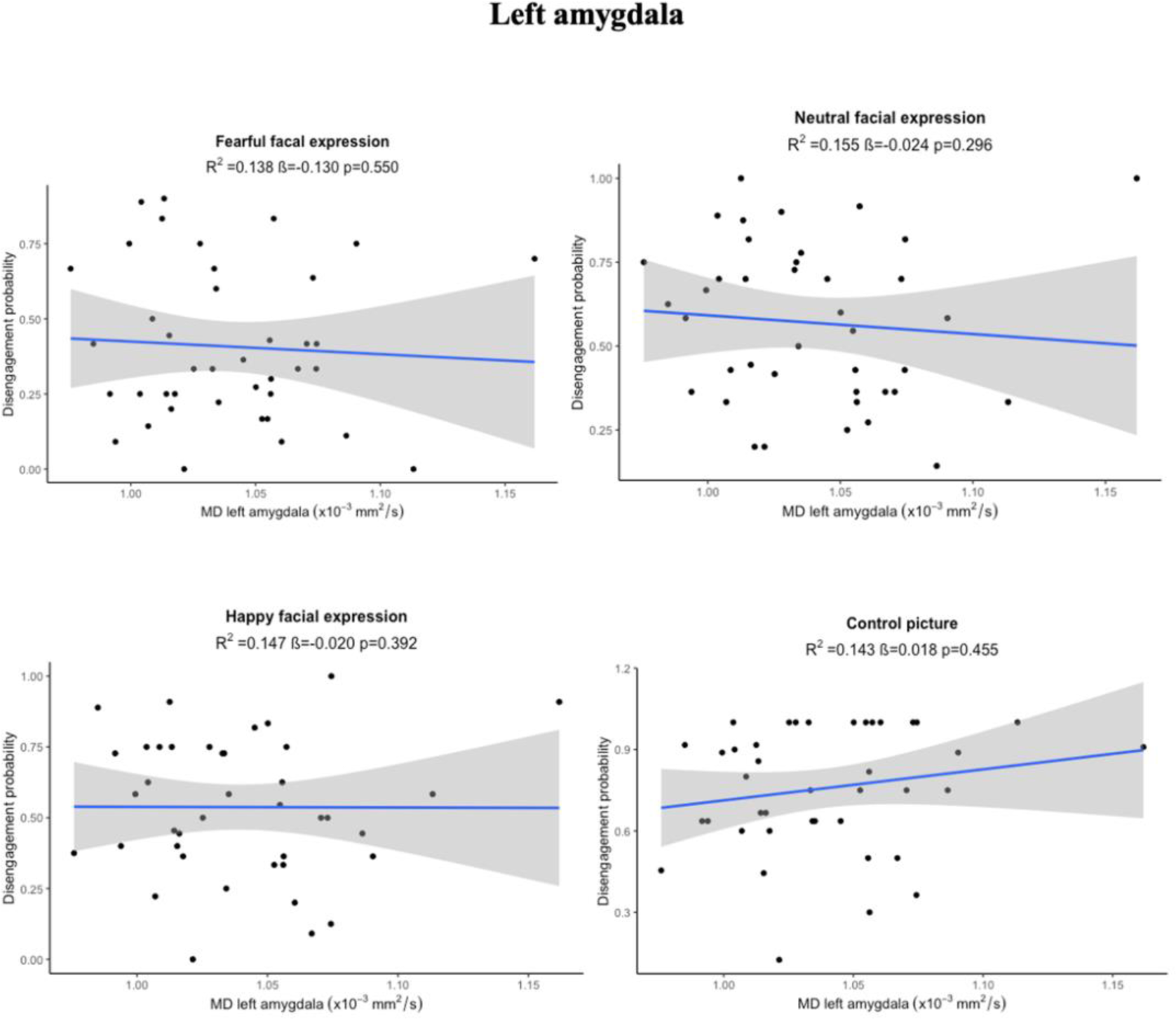
The disengagement probability from fearful, non-fearful (happy and neutral), and scrambled non-face control pictures in the overall sample.

**Table 3.** Multiple linear regression model for the main and sex-interaction associations for the disengagement probability (DP) from fearful (FE), happy (HA), neutral (NE) and scrambled non-face control picture (CS) faces.

|  | Left amygdala MD |  |  | Right amygdala MD |  |  |
| --- | --- | --- | --- | --- | --- | --- |
| | $\beta$ (P) | SE | $R^2$ | $\beta$ (P) | SE | $R^2$ |
| **Main association** |  |  |  |  |  |  |
| DP FE+ age at the MRI scan date | -.01(.55) | .02 | .13 | -.05(.04) | .02 | .06 |
| DP Ha + age at the MRI scan date | -.02(.39) | .02 | .14 | -.04(.20) | .03 | .0004 |
| DP NE + age at the MRI scan date | -.02(.29) | .02 | .15 | -.06(.05) | .03 | .05 |
| DP CS + age at the MRI scan date | .01(.45) | .02 | .14 | .02(.65) | .001 | -.03 |
| **Sex-interaction association** |  |  |  |  |  |  |
| sex * DP FE+ age at the MRI scan date | .02(.59) | .04 | .15 | .004(.94) | .05 | .04 |
| sex * DP HA + age at the MRI scan date | -.07(.11) | .04 | .21 | -.09(.14) | .06 | .04 |
| sex * DP NE + age at the MRI scan date | -.01(.79) | .04 | .16 | -.04(.47) | .06 | .05 |
| sex * DP C + age at the MRI scan date | -.11(.019) | .04 | .27 | -.19(.004) | .06 | .16 |

The sex-interaction analyses revealed no significant associations between the amygdala MD measures and the attentional disengagement probability from fearful, non-fearful (neutral and happy) faces. However, there was a significant sex-interaction association between the MD of the right and left amygdala and the disengagement probability from the scrambled non-face control picture after controlling for infant age at the time of the MRI scans (for the right amygdala MD, *ß* = -.19, *p =* .004 and for the left amygdala MD, *ß* = -.11, *p =* .019). The higher the right and left amygdala MD, the higher the disengagement probability from scrambled non-face control picture in girls (for the right amygdala MD, *ß* =.130, *p =* .021 and for the left amygdala MD, *ß* =.19, *p =* .013) (Figure 3) (Table 3). In boys, there was no significant relationship between amygdala mean diffusivity and disengagement probability from fearful and non-fearful (neutral and happy) faces.

**Figure 3.**
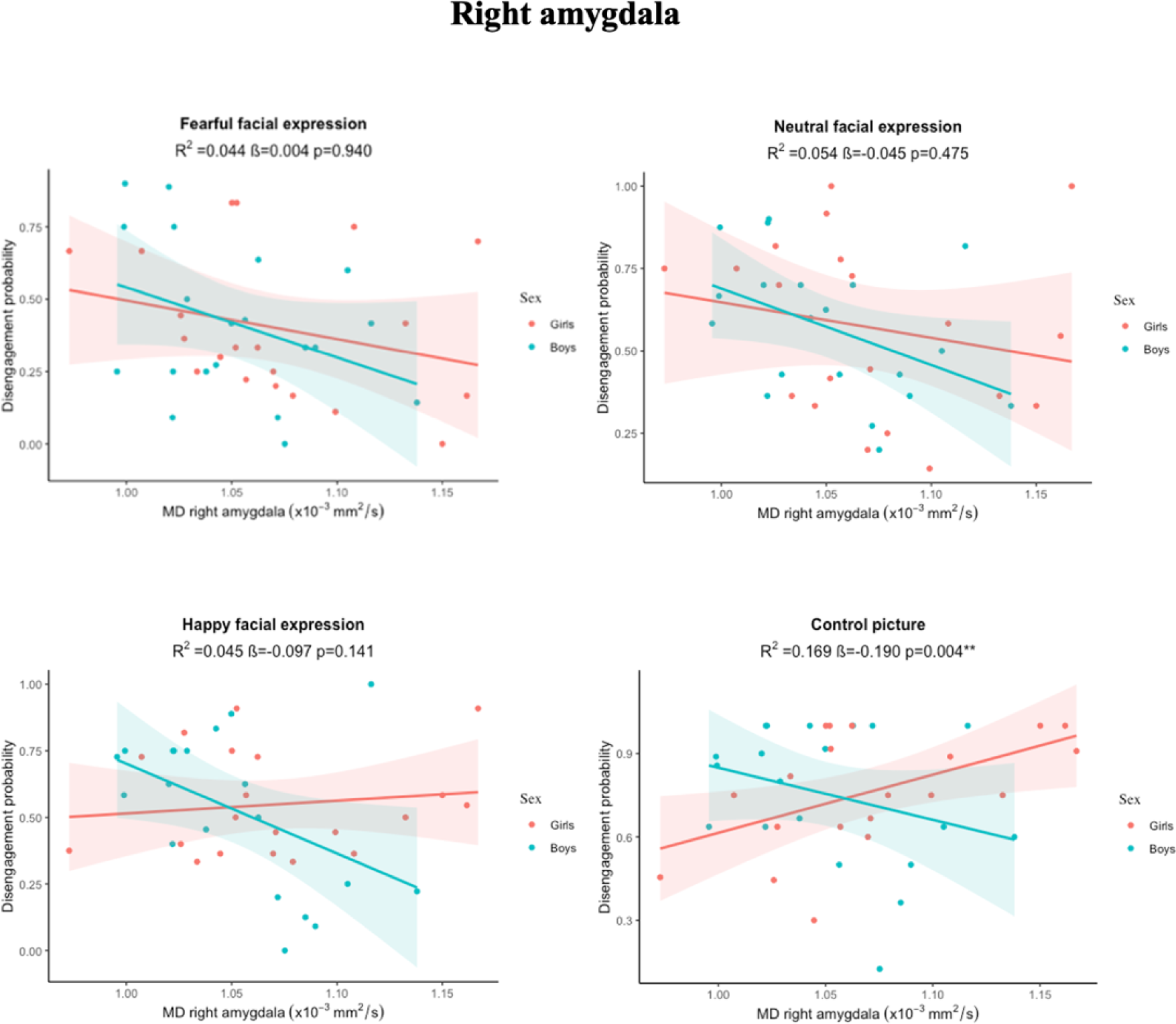
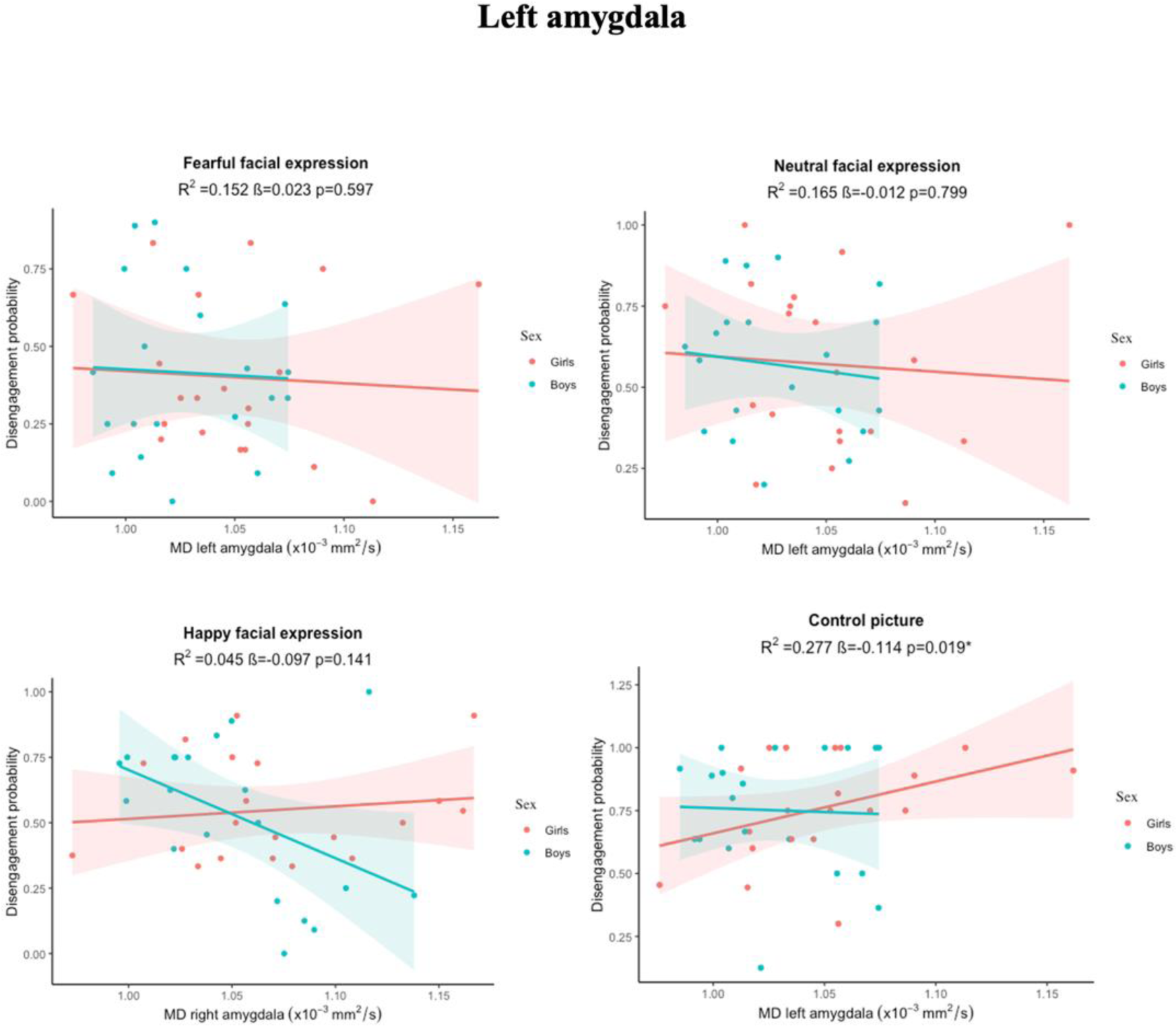
Right and left amygdala mean diffusivity (MD) values and disengagement probability from fearful, non-fearful (neutral and happy), and scrambled non-face control pictures in girls and boys.

## Discussion

In this study, we revealed for the first-time significant associations between the amygdala MD and the disengagement probability from fearful faces in infants. These results showed that the higher the right amygdala MD measure, the lower the probability of disengaging attention from fearful faces vs. non-fearful faces (neutral and happy) towards salient lateral distractors in infants. Our findings support earlier studies in infants that show the amygdala’s role in attentional bias in processing fearful facial expressions when compared to other facial expressions (Tuulari et al., 2020; Vuilleumier, 2005; Whalen et al., 2001). During the embryonic stage, the amygdala, one of the first structures to emerge, experiences a significant acceleration in its developmental process (Acosta et al., 2019; Humphrey, 1968; Müller & O’Rahilly, 2006). Subsequently, during infancy, the amygdala continues to rapidly mature and becomes increasingly responsive to emotional stimuli (Tottenham & Laurel J. Gabard-Durnam, 2019; Tottenham & Sheridan, 2010; Uematsu et al., 2012). While the specific neural mechanisms that direct infants’ attention to fearful faces are unknown, current evidence indicates that the amygdala and orbitofrontal cortex are involved in the processing of all types of facial expressions, not just fearful ones (Fitzgerald et al., 2006). Consequently, the role of the amygdala in the perception and recognition of emotions appears to be more modulatory and temporally extended (Adolphs, 2010). Overall, the infant amygdala’s sensitivity to emotional stimuli is an important aspect of early emotional and social development (Leppanen & Nelson, 2010; Tottenham, 2013).

The results from this study might suggest that changes in the microstructural properties of the amygdala, such as the MD measure, may contribute to the processing of fear-related information. Higher MD in the right amygdala may indicate altered microstructures potentially impacting fear processing (Salzwedel et al., 2019; Yrondi et al., 2019). Infants with elevated MD levels may show heightened sensitivity to fearful faces, leading to reduced attentional disengagement from these stimuli. This heightened sensitivity could suggest an attentional bias towards threat, maintaining focus on fearful faces (Fu & Pérez-Edgar, 2019; Koelkebeck et al., 2021), thereby decreasing attentional shifts toward other salients. Differences in amygdala microstructure may reflect variations in early neurodevelopment (Yrondi et al., 2019), potentially leading to atypical neural maturation and altered attentional patterns. Interactions between the amygdala and other brain regions involved in attentional control and salience processing could further influence attentional processing, impacting the probability of disengaging attention from fearful faces towards other stimuli. Additionally, infants with higher MD in the right amygdala may have underlying biological vulnerabilities affecting attentional processes, particularly in response to fear-related stimuli (He et al., 2017), potentially manifesting as reduced attentional disengagement. It’s worth noting that these differences in amygdala MD can potentially become more comparable to typical patterns with increasing age. Moreover, higher MD measures in the amygdala may indicate an early neurobiological vulnerability to fear dysregulation, which may manifest as difficulty disengaging from signals of a potential threat in infancy. Previous studies have indicated that there are multiple physiological mechanisms that may be responsible for changes in MD. Increased MD refers to an increase in the rate of water molecule diffusion (Cherubini et al., 2010). Increased MD may indicate a disruption in the microstructural integrity of the tissues, increased extracellular spaces, as well as shrinkage of neurons, and synapses (Cherubini et al., 2010; De Gennaro et al., 2011; Gillespie et al., 2017; Kantarci et al., 2005). In patients with anxiety, traumatic brain injury, cognitive impairment, and Alzheimer’s disease, increased MD in the amygdala has been linked to reduced microstructural integrity (Cherubini et al., 2010; Juranek et al., 2012), but also with higher empathizing, cooperativeness, and cognitive ability (Takeuchi et al., 2019). Reduced MD indicates cytoarchitectonic changes and increased neural tissue density (Beaulieu, 2002; Sagi et al., 2012). Moreover, a decrease in the amygdala MD has been observed after electroconvulsive therapy in patients with treatment-resistant depression, indicating a normalization of tissue integrity (Yrondi et al., 2019). The presented results add to the previous research suggesting that early changes in the microstructural properties of the amygdala, such as the MD measure, may contribute to the processing of fear-related information in infants. Since a decrease in MD measures has been observed throughout childhood and adolescence, it is commonly considered that lower MD measures correspond to a more mature brain and higher functional adaptation (Takeuchi et al., 2016; Tamnes et al., 2010; Yoshida et al., 2013). Furthermore, while changes in amygdala MD measures may be associated with negative outcomes in offspring, decreased MD is not always associated with increased cognitive ability or functional adaptation (Takeuchi et al., 2015). However, the interpretation of changes in the amygdala MD in infants can be complex, and the underlying physiological mechanisms and implications need further investigation.

The sex-interaction analyses in our study revealed that girls had significantly stronger associations between the probability of disengaging from scrambled non-face control picture and the MD of both the right and left amygdala in comparison to boys. The observed association between higher MD in the amygdala and quicker attentional disengagement from non-emotional control pictures towards distractors in girls suggests that elevated MD levels may correspond to more efficient attentional networks facilitating rapid redirection of attention away from non-salient stimuli. The precise mechanisms underlying these observed results are unknown. Research has shown different patterns of sex differences in amygdala responses to faces and emotional stimuli (Killgore et al., 2001). Women have been found to have more bilateral amygdala activity to emotional stimuli than men (Hamann, 2005; Killgore et al., 2001). While some studies have suggested that sex differences in amygdala responses to faces may play a role in women’s enhanced emotional reactivity (Hall & Matsumoto, 2004), the evidence is not conclusive. Other studies have found the opposite pattern, with men showing greater amygdala activation in response to emotional stimuli (Derntl et al., 2009; Kret et al., 2011). These findings suggest that there is considerable variability in amygdala responses to emotional stimuli, more research is needed to fully understand the underlying mechanisms of these sex differences in response to emotional faces.

Studies have demonstrated that there is a clear difference in how the left and right amygdalae process information, as evidenced by lateralized amygdala activity in each hemisphere (Baas et al., 2004). While the right amygdala is primarily responsible for the rapid and initial identification of stimuli, the left amygdala may play a greater role in the subsequent processing and interpretation of the identified stimuli, with a slower response time (Bonnet et al., 2015; Sergerie et al., 2008). Several studies have revealed that fearful faces stimulate the right amygdala more than the left (Framorando et al., 2021; Liddell et al., 2005; Morris et al., 1999). Other studies, on the other hand, have found that the left amygdala regulates attentional reactions to fearful faces (Carlson et al., 2009; Thomas et al., 2000; Tuulari et al., 2020). The right amygdala’s response to fearful faces could be due to the right amygdala’s stronger connection to visual processing areas in the brain (Markowitsch, 1998; Xiao et al., 2021). This connection helps to recognize and respond to threatening or fearful stimuli more rapidly and effectively, resulting in a heightened response of the extrastriate cortex via fast amygdalofugal projections to the visual regions (Framorando et al., 2021). However, the precise role of lateralization in amygdala activation remains unknown (Baas et al., 2004). More research is needed to investigate the neural mechanisms and developmental significance of the associations between amygdala diffusivity and attentional bias toward fearful faces, particularly in terms of hemispheric specificity.

A few limitations of this work should be acknowledged. DTI is frequently used to reveal the microstructure of white matter brain structures (Salat, 2013). The analysis of gray matter (GM) using DTI is currently constrained by several factors. These include the limited presence of GM in certain brain regions such as the amygdala, the overlapping partial volume effect with the small amount of GM, and the low resolution of DTI (Salat, 2013). Moreover, MD might be the only measure suitable to be measured in GM structures as, unlike in white matter, the diffusivity in GM is isotropic (Gillespie et al., 2017; Jeurissen et al., 2013; Sexton et al., 2010; Yrondi et al., 2019). Alternatively, advancements in the accuracy and sensitivity of DTI in GM structures are needed (Salat, 2013). In this study, we used a high diffusion direction (60) to achieve better tensor model fitting and results (Hashempour et al., 2023; Tian et al., 2020). Due to the lower image resolution and movement artifacts, this is particularly useful for imaging infant brains (Gousias et al., 2012; Hashempour et al., 2023; Weisenfeld et al., 2006). While our sample size was small it contained a higher level of variation. Furthermore, due to the relatively small sample size in our study, we recommend that future research aim to replicate these findings while addressing the aforementioned limitations and incorporating a larger sample size.

## Conclusion

In conclusion, we demonstrated for the first time that the MD of the amygdala is related to attention disengagement processes already in infancy, both in threat processing (in the whole sample, higher amygdala MD was associated with more delayed attention disengagement from fearful faces towards distractors) and in non-emotional conditions (in infant girls, higher amygdala MD was associated with a more frequent attention disengagement from non-faces towards distractors). Our results also show that the amygdala MD properties are differently associated with attention disengagement tendencies in scrambled non-face control picture and fearful face conditions. Future research may benefit from investigating the biological pathways that underpin these associations, as well as how the observed amygdala diffusivity changes predict offspring emotional and social behaviors.

## Acknowledgments and disclosures

NH was supported by the University of Turku graduate school. JJT was supported by Sigrid Jusélius Foundation; Emil Aaltonen Foundation; Finnish Medical Foundation; Alfred Kordelin Foundation; Juho Vainio Foundation; Turku University Foundation; Hospital District of Southwest Finland; State Grants for Clinical Research (ERVA); Orion Research Foundation, Signe and Ane Gyllenberg Foundation. HM was supported by Academy of Finland (#26080983). NMS was supported by the Hospital District of Southwest Finland; State Grants for Clinical Research (ERVA). RK was supported by Ane & Signe Gyllenberg Foundation ja Academy of Finland (308252). LK was supported by the Academy of Finland (#325292/Profi5, #308176), Signe and Ane Gyllenberg Foundation, State Grants for Clinical Research (ERVA), NARSAD Brain and Behavior Research Foundation YI Grant #1956. HK was supported by Finnish Academy No. 264363, 253270, 134950; and Signe and Ane Gyllenberg Foundation and Jane and Aatos Erkko Foundation. ELK was supported by Finnish Cultural Foundation and Alli Paasikivi Foundation.

We would like to thank Kristiina Kuvaja for carrying out the scans with the investigators and especially all FinnBrain families that participated to the measurements. We thank Dr. Jukka Leppänen for his contribution to the eye-tracking data analyses.

The authors declare no competing interest.

